# Isolation and characterization of the new *Streptomyces* phages Kamino, Geonosis, Abafar and Scarif infecting a broad range of host species

**DOI:** 10.1101/2024.02.26.582091

**Authors:** Bente Rackow, Clara Rolland, Isabelle Mohnen, Johannes Wittmann, Mathias Müsken, Jörg Overmann, Julia Frunzke

## Abstract

*Streptomyces*, a multifaceted genus of soil-dwelling bacteria within the phylum Actinobacteria, features intricate phage-host interactions shaped by its complex life cycle and the synthesis of a diverse array of specialized metabolites. Here, we describe the isolation and characterization of four novel *Streptomyces* phages infecting a variety of different host species. While phage Kamino, isolated on *Streptomyces kasugaensis*, is predicted to be temperate and encodes a serine integrase in its genome, phages Geonosis (isolated on *S. griseus*) and phages Abafar and Scarif, isolated on *S. albidoflavus*, are virulent phages. Phages Kamino and Geonosis were shown to amplify well in liquid culture leading to a pronounced culture collapse already at low titres. Determination of the host range by testing >40 different *Streptomyces* species identified phages Kamino, Abafar and Scarif as broad host range phages, with Kamino showing productive infections on 15 different species. Isolated phages were further tested regarding their sensitivity to antiphage molecules. Here, the strongest effects were observed for the DNA-intercalating molecule daunorubicin. Overall, the phages described in this study expand the publicly available portfolio of phages infecting *Streptomyces* and will be instrumental for advancing the mechanistic understanding of the intricate antiviral strategies employed by these multicellular bacteria.

## Introduction

*Streptomyces*, a genus of soil-dwelling multicellular Gram-positive bacteria belonging to the phylum *Actinomycetota*, stands out for its extensive array of biosynthetic gene clusters. These clusters encode a wide range of specialized metabolites exhibiting diverse biological activities, such as antibacterial, antifungal, anticancer, and even antiviral properties. Today, up to two-thirds of all nature-based antibiotics used in the clinics are produced by *Streptomyces* spp ^1–3^. Furthermore, the intricate life cycle of *Streptomyces* positions it as a model for the study of multicellular development in bacteria ^4^. Microbial interaction was shown to be a prominent trigger of secondary metabolite production and cellular development. Co-cultivations of microorganisms therefore represent a promising approach for the discovery of novel antibiotics otherwise not produced under laboratory conditions ^5^. In recent years, also the interaction of *Streptomyces* with their most abundant predator in the environment – bacteriophages (or phages for short) – increasingly gained attention ^6,7^.

Phage research in previous decades focused on phages for tool development employing phages like phiC31 or R4 ^8–10^. Genetic tools such as restriction enzymes and integrative plasmids have been constructed and are still valuable tools used in *Streptomyces* genetics ^11,12^. In recent years, however, the focus of actinobacteriophage research has shifted from tool design to phage-host interaction, in particular to the elucidation of novel antiviral defence mechanisms ^13^. Remarkably, it was shown that secondary metabolites produced by *Streptomyces*, belonging to the classes of anthracyclines and aminoglycosides, inhibit infection by a broad range of phages ^14,15^. Furthermore, cellular development was shown to play a key role in the emergence of transient resistance to phage infection. This was shown for the development of transiently resistant mycelium at the infection interface on plates ^16^ as well as for the formation of S-forms lacking the cell wall, which occurred during infections under osmoprotective conditions ^17^.

For the discovery and mechanistic understanding of novel defense strategies a diverse set of phages is needed incentivising the isolation and characterization of new phages. Using different *Streptomyces* species as isolation host, several new phages have been isolated and described in recent years ^18–20^. These studies show that the common notion of a narrow host range and specific infection of phages does not necessarily apply to *Streptomyces* phages, as several actinophages have been described to have a broader host range and to productively infect several species of the genus ^20,21^. However, in most studies only a small number of different host species has been tested delimitating the information regarding host range.

In this study, we isolated and characterized four new phages infecting *Streptomyces*. Testing a collection of over 40 distinct *Streptomyces* strains, we observed that the phages Kamino, Abafar, and Scarif exhibited the ability to infect a wide array of *Streptomyces* species. This characteristic emphasizes their suitability for comparative studies on phage defense mechanisms across various host species. All four phages isolated in this study were sequenced and characterized regarding their infection dynamics and their sensitivity to known antiphage compounds.

## Materials and methods

### Bacterial strains and growth conditions

*Streptomyces albidoflavus* M145, *Streptomyces griseus* and *Streptomyces kasugaensis* were used as main host strains in this study. Bacterial cultures were started from a spore stock, stored at – 20°C in 20 % glycerol. Spores were inoculated into fresh liquid Glucose Yeast Malt extract (GYM) medium for *S. griseus* and *S. kasugaensis* or into Yeast extract Malt extract (YEME) medium for *S. albidoflavus* M145. In general, cultivation was carried out at 30 °C. Double agar overlays were performed on GYM agar for all species, were 4 ml 0.4 % agar and 20 ml 1.5 % agar was used for the top and bottom layer, respectively.

### Phage isolation and propagation

Phages described in this study were isolated from soil samples taken in the Eifel region (Aachen, Germany) and Braunschweig, respectively. The soil samples were incubated with sodium chloride / magnesium-sulphate (SM) buffer (10 mM Tris-HCl pH 7.3, 100 mM NaCl, 10 mM MgSO_4_, 2 mM CaCl_2_) over night at 4 °C on a shaking plate, to resolve phages from soil particles ^18^. The samples were then centrifuged for 30 min at 5000 x *g* to separate the supernatant from soil particles. The supernatant was subsequently filtered through a 0.22 µm pore size membrane filter, to remove remaining bacteria. The lysate was furthermore enriched on the host bacteria described above, to enrich potential phages overnight. After overnight incubation serial dilutions were spotted on a bacterial lawn propagated by mixing overnight cultures of the host strains with 4 ml top agar to a final OD_450_= 0.4. Plaques were visualized after incubation over night at 30 °C.

Pure phages were obtained by restreaking single plaques twice on a double agar overlay. Amplification of phages for high titre stocks was performed by mixing 100 µl phage sample with the top agar of a double agar overlay to obtain confluent lysis of the bacterial lawn. After overnight incubation at 30 °C, the phages were resolved by adding 5 ml SM buffer to the plates and shaking the agar plates for 1-2 hours at room temperature at 40 rpm. The SM buffer was then collected from the plates, centrifuged and subsequently filtered through a 0.22 µm membrane filter to separate remaining bacteria from the phage lysate. The high titre lysate was mixed with 10 % (v/v) sterile glycerol and stored at either −80 °C for long-term storage or 4 °C for further experiments. The phage titre of the lysate was determined by spotting 2 µl of a serial dilution up to 10^-8^ onto a double agar overlay with the top agar containing the host bacterium to an OD_450_ = 0.4. After overnight incubation at 30 °C, the highest dilution with visible single plaques was counted and the titre in plaque forming units (PFU) per ml was calculated.

### Electron microscopy

For transmission electron microscopy (TEM) 5 µl of pure high titre phage lysate were dropped on a glow discharged (15mA, 30 s) carbon-coated copper grid (CF300-CU, carbon film 300 mesh copper). The phage containing grid was stained with 2 % (w/v) uranyl acetate for 5 minutes and washed twice in ddH_2_O. Dried samples of Kamino and Geonosis were analysed with a TEM Talos L120C (Thermo Scientific, Dreieich, Germany) at an acceleration of 120 kV. Abafar and Scarif samples were examined in a Zeiss EM 910 or Zeiss Libra120 Plus transmission electron microscope (Carl Zeiss, Oberkochen, Germany) at an acceleration voltage of 80 kV/120 kV at calibrated magnifications using 300 mesh copper grids and a mica-floated carbon film enabling attachment of phages.

### Infection dynamics

Infection experiments in liquid cultures were performed as described in Hardy et al. (2020). Cultivation was performed in the BioLector micro cultivation system of m2plabs (Aachen, Germany) as biological triplicates in 48-well flower plates (m2plabs) at 30 °C with a shaking frequency of 1200 rpm. Backscattered light intensity was measured every 15 minutes (filter module: e_xcitation_/e_mission_, 620 nm/ 620 nm; gain 25) and supernatant samples were taken every 2 hours to assess the infection dynamics and amplification rate. Infection took place in 1 ml YEME or GYM medium with *S. albidoflavus* M145*, S. griseus* and *S. kasugaensis* with an initial OD_450_ of 0.15. Phages were directly added to the cells with initial titres from 10^2^ to 10^7^ PFU/ml. The sampled supernatant was centrifuged to separate phages from cell remnants and subsequently diluted in a serial dilution of 10^-1^ to 10^-8^. The dilutions were spotted on a GYM double agar overlay containing the respective host bacteria in the top agar layer with an OD_450_ = 0.4 to determine the phage titre over the course of the infection.

### Plaque development

The plaque morphology of the phages Abafar, Geonosis, Kamino and Scarif were observed on double agar overlay plates with a titre of 10^2^ to 10^3^ PFU/ml of each phage and an OD_450_ = 0.4 of the respective host bacterium in the top agar. The plaque assay plates were incubated at 30 °C for 72 hours in total and images of single plaques were taken using a Nikon SMZ18 stereomicroscope with NIS-Elements AR 5.3 software after 24 h, 48 h and 72 h.

### Phage host range assay

Host range determination of the four phages was performed with classical spot assays as described in the section “Phage isolation and propagation”. Different *Streptomyces* species (see Table S1 for productive infections and Table S2 for all tested strains) were used as bacterial lawn to test the infectivity of the respective phages. In this study, we discriminated between the simple clearance of the bacterial lawn by the phages and truly productive infections, where single plaques of the phages could be observed on the bacterial lawn. For all productive infections the efficiency of plating (EOP) was calculated by dividing the counted plaques on the tested bacterial host by the counted plaques on the “original” host, which was used for isolation.

### Sensitivity to antiphage molecules

All phages were tested for their sensitivity towards small molecules, which are known to have anti-phage properties, such as aminoglycosides (apramycin and hygromycin), anthracyclines (daunorubicin) and the DNA-intercalating peptide antibiotic actinomycin D ^22–24^. To determine whether the phages showed sensitivity towards the anti-phage compounds, spot assays were carried out as described before, however in this case rising concentrations of the antibiotics were added to the top and bottom layers of the double agar overlay. The spot assays were carried out in duplicates and all four phages were tested against daunorubicin and actinomycin D at the concentrations 0 µM, 1 µM, 3 µM and 6 µM and against apramycin and hygromycin at the concentrations 0 µg/ml, 2.5 µg/ml, 25 µg/ml and 50 µg/ml. The final titre was determined after 24-48 hours of incubation at 30 °C.

### DNA isolation, genome assembly and annotation

Genomic DNA of the phages was isolated from 1 ml phage lysate with a high titre using the NORGEN BIOTEK CORP. Phage DNA isolation Kit (Thorhold, Canada). Isolation was carried out as described in the manual provided by the manufacturer, including all optional steps. To increase the concentration a two-step elution of the DNA in each 25 µl elution buffer was performed. DNA concentration was measured in a NanoPhotometer (P330, IMPLEN, Germany). Whole genome sequencing using the Illumina NovaSeq platform with a read length of 2x 150 bp was performed by GENEWIZ Germany. The NEBNext Ultra II DNA Library Prep Kit was used to sequence the whole genome of phages on an Illumina MiSeq platform with paired reads of 15-150 base pairs in length. Initially, quality control checks for each pair of raw sequencing reads was performed using FASTQC v0.11.9 (http://www.bioinformatics.babraham.ac.uk/projects/fastqc/). The adapter and low-quality reads were cleaned and trimmed from the sequencing data with the help of the fastp v0.23.2 program ^25^. Next, the whole genome *de novo* assembly with the trimmed high-quality reads, was performed with the help of the Shovill pipeline v1.1.0 (https://github.com/tseemann/shovill) using the SPAdes genome assembler v3.15.5 ^26^. Lastly, Pilon version 1.24 ^27^ was used to improve, and curate the assembled genomes. Phage genomic terminal ends were identified using the PhageTerm ^28^ online galaxy platform.

All phage genomes were annotated with Prokka 1.8 using different databases (Markov model profile databases, including Pfam and TIGRFAMs). A search was performed using hmmscan from the HMMER 3.1 package ^29^. Sequences were also compared to the PHROG database ^30^. Taxonomic classification was determined on the identity level to close related phages using NCBI BLASTn (NCBI, 2023) search and ordered according to the new virus taxonomy release (ICTV, 2023).

## Results

### Bacteriophage isolation and morphology

Four novel phages infecting different *Streptomyces* species were isolated from soil. Phage Kamino was isolated on *S. kasugaensi*s as host, Geonosis on *S. griseus*, as well as Abafar and Scarif on *S. albidoflavus* (previously *S. coelicolor*) as isolation host. Phage Kamino forms small, turbid plaques (Figure 1A) with a plaque area of approximately 0.3 mm^2^; the plaque size is relatively constant over 48 hours of incubation (Figure 1B). Phage Geonosis forms round and clear plaques with sharp edges (Figure 1 A) and a plaque area of approximately 4 mm^2^ after 24 h of incubation, which increases up to 38 mm^2^ on average after 72 hours of incubation (Figure 1B). The *S. albidoflavus* infecting phages Abafar and Scarif show an overall similar plaque size as Geonosis with average plaque areas of 4.2 and 4.0 mm^2^, respectively, after 24 hours incubation. Abafar and Scarif also show an increase in plaque area over the course of 72 hours up to an average plaque area of 17 and 21 mm^2^ (Figure 1B). Additionally, around the plaques of Abafar and Scarif enhanced production of actinorhodin was observed by the formation of coloured halos around the plaques, which has been described in previous studies reporting *S. albidoflavus/S. coelicolor* phages ^18,19^(Figure 1A).

**Figure 1:**
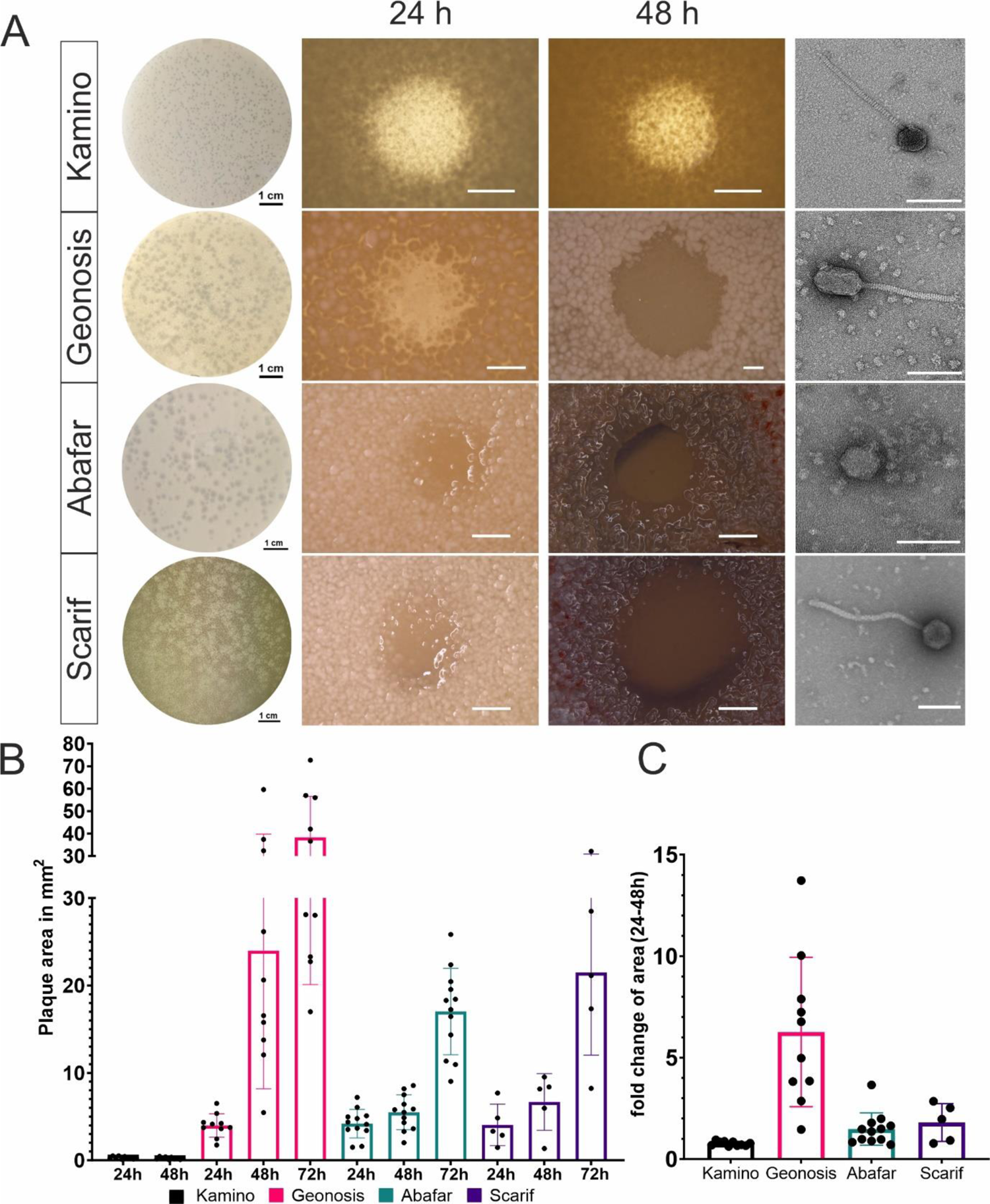
Morphological comparison of plaques and virions. A) Plaque and TEM images of the tested phages with an overview image of a plate with plaques, a plaque close up image 24 h and 48 h post infection and a TEM image of the virion from left to right respectively for the phages Kamino, Geonosis, Abafar and Scarif from top to bottom. The scale bar for the overview images is 1 cm in length, for plaque close ups the scale bar is 1000 µm and for TEM images 100 nm. B) Comparison of the development of plaque area over time of 24-72 hours post infection with plaque area shown in mm^2^ for the phages Kamino (black, n= 10) Geonosis (pink, n= 10), Abafar (turquoise, n= 12) and Scarif (purple, n= 5). C) Fold change of plaque area between 24 h and 48 h post infection for all four phages in the same colour as graph B.

TEM imaging of the phage particles revealed that phages Kamino, Geonosis and Scarif belong to the morphotype siphovirus with an icosahedral capsid and a long, non-contractile tail, whereas phage Abafar features a podoviral morphotype, showing an icosahedral capsid with a very short tail (Figure 1 A right column). Details on capsid and tail length are provided in Table 1.

**Table 1:** Morphological comparison of virions.

| Phage | Tail length [nm] | Capsid length [nm] | Capsid diameter [nm] |
| --- | --- | --- | --- |
| **Kamino (n=13)** | 223.4 (+/- 7.1) | 63.5 (+/- 2) | 61.5 (+/- 1.4) |
| **Geonosis (n=15)** | 176.5 (+/- 4.4) | 88.6 (+/- 4.8) | 53.8 (+/-3.0) |
| **Abafar (n=8)** | 15.7 (+/- 1.5) | 63 (+/- 3.1) | 63 (+/- 1.9) |
| **Scarif (n=5)** | 270.2 (+/- 4.9) | 69.8 (+/- 1.9) | 67.4 (+/- 1.4) |

### Infection curves and host-range of bacteriophages

In order to assess phage infection dynamics, phage infections in liquid cultures on the original host strain were performed. As it is not suitable to perform one-step growth curves with *Streptomyces* spp. due to the multicellular development, the host strains were cultivated in microtiter plates in presence of different phage titres. Cell growth was monitored by backscatter measurements in 15 minutes intervals over the course of 24 hours and phage propagation was determined by taking samples from the culture supernatants every 2 hours to determine the titre of infectious phage particles at the respective time (Figure 2). Infections with phage Kamino and phage Geonosis lead to a complete culture collapse of their respective hosts, *S. kasugaensis* and *S. griseus,* with a starting titre as low as 10^2^ PFU/ml. The titre of Kamino and Geonosis increased steadily and reached final titres of 10^6^ to 10^9^ PFU/ml for Kamino and 10^10^ to 10^11^ PFU/ml for Geonosis. On high titres (10^7^), however, little to no amplification was observed for phage Kamino. In contrast, phages Abafar and Scarif showed little to no growth defect on the bacterial culture, independent of the starting titre. Amplification of Abafar and Scarif could only be observed in spot assays, determining the phage titre over time. In case of phage Abafar, amplification in liquid was first observed for a starting titre of 10^4^ PFU/ml and for phage Scarif at a starting titre of 10^5^ PFU/ml, which indicates a probably lower burst size and decreased infectivity compared to Geonosis and Kamino. While the final titre determined for phages Kamino and Geonosis appeared independent from the starting titre, final titres significantly increased for Abafar and Scarif, when infection was initiated with higher starting titres. Under the specified conditions, Abafar achieved final titres of 10^6^ to 10^12^ PFU/ml 24 h post infection while Scarif achieved titres between 10^4^ to 10^6^ PFU/ml. Altogether, all four phages are able to propagate in liquid cultures to different extends. Abafar and Scarif however reach higher titres when the lysate is prepared on plates.

**Figure 2:**
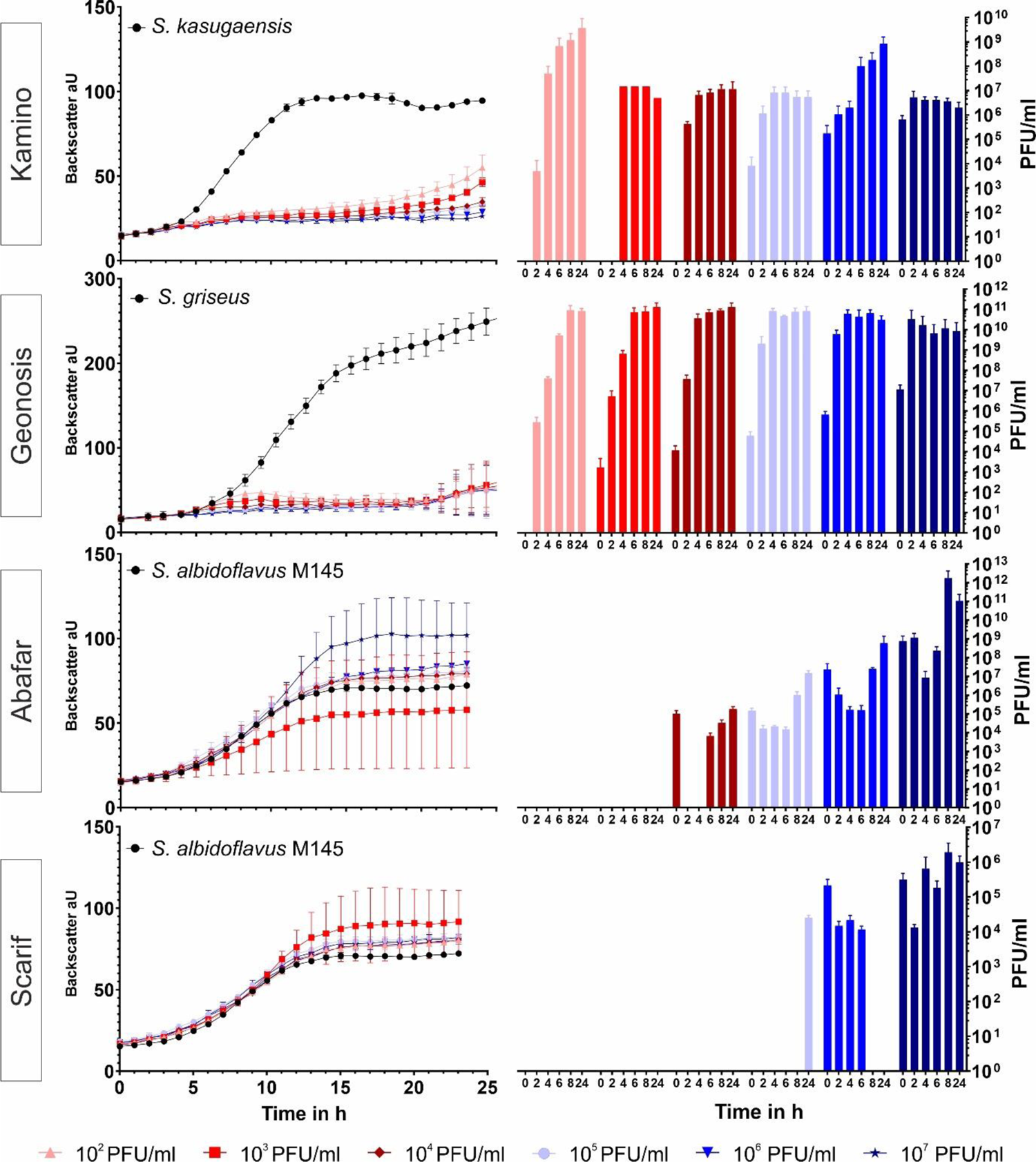
Infection curves of the four phages on their isolation host *S. kasugaensis* infected by Kamino, *S. griseus* infected by Geonosis and *S. albidoflavus* M145 infected by Abafar and Scarif. *S. kasugaensis* and *S. griseus* were inoculated to GYM medium and *S. albidoflavus* was grown in YEME medium. All strains were cultivated in microtiter plates and infected with increasing titres of the respective phages. In the left panel, the backscatter is plotted against time to visualize growth of the bacterial culture. In the right panel phage titre at different time points throughout the infection (2/4/6/8 and 24 h) are shown. The colours of the growth curves and the bar plots for the phage titre correlate to the same initial infection titre between 10^2^ to 10^7^ PFU/ml, the black curve indicates growth of the host bacterium in absence of phages. n= 3 independent biological replicates.

### Host Range of phage isolates

One physiological trait that is important to consider when using or studying phages, is the range of bacterial hosts they are able to infect. In this study, we determined the host ranges of the novel phage isolates by spotting serial dilutions of the phages on lawns of >40 different *Streptomyces* species (Table S1). A distinction must be made between the simple lysis of bacteria and the ability to cause a productive infection, because only the latter leads to the formation of individual plaques (Figure 3B). From the four phages described in this study, phage Kamino has the broadest host range with lawn clearance on 22 *Streptomyces* spp. and productive infections on 15 different species among the 45 species tested (Table S1, Figure 3B). The efficiency of plating (EOP) refers to the ratio of plaques formed on the host used for isolation compared to another host species. While phage Kamino is able to infect a wide variety of host strains, the EOP ranges from 0.002 % on *S. olivaceus* up to 5000 % on *S. afghaniensis* (Figure 3A). The phages Abafar and Scarif have the same range of productive infections with 7 different species but differ in their lawn clearance, where Abafar shows clearance on 12 species and Scarif only on 10 (Figure 3B). Phage Abafar displays an EOP lower than 100 % on the alternative hosts. Scarif however, reaches EOP’s up to 8000 % on *S. afghaniensis* (Figure 3A). In contrast to these broad host-range phages, phage Geonosis is only able to infect two different *Streptomyces* species, its isolation host *S. griseus* and one additional host, *S. olivaceus* with an EOP of 0.15 %, which classifies Geonosis as a narrow host-range phage in this context (Figure 3A, B).

**Figure 3:**
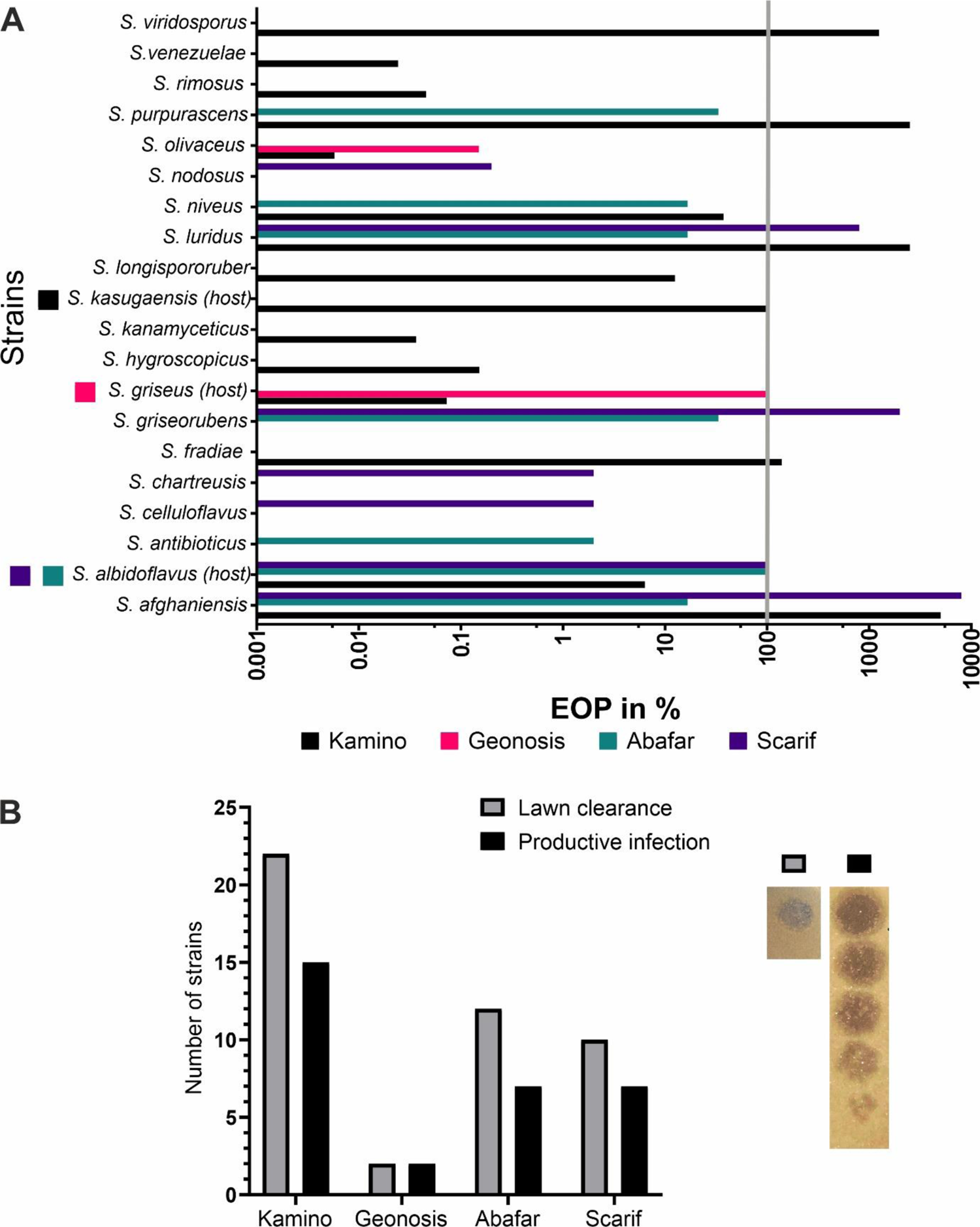
Host range of the *Streptomyces* phages Kamino, Geonosis, Abafar and Scarif. A) Efficiency of plating (EOP) in % to describe the host range of the four phages. On the Y-axis the tested *Streptomyces* strains are shown and on the X-axis the EOP in %. 100 % EOP (obtained on the isolation host) is marked with the grey line. B) Distribution of productive infection (black) versus lawn clearance (grey). On the right are exemplary images of lawn clearance and productive infection of phage Scarif on *S. albidoflavus*.

### Comparison of genomes

Phage DNA of all four phages was isolated and sequenced using Illumina short read sequencing. Reads were assembled and contigs were annotated using Prokka with implemented PHROG analysis (Table 2). Prediction of the phage lifestyle was performed with the machine-learning tool PhageAI ^31^ with 93.8 % confidence for a temperate lifestyle of Kamino and 93.5 %, 95.0 % and 73.6 % confidence for Geonosis, Abafar and Scarif having a virulent lifestyle, respectively.

**Table 2:**
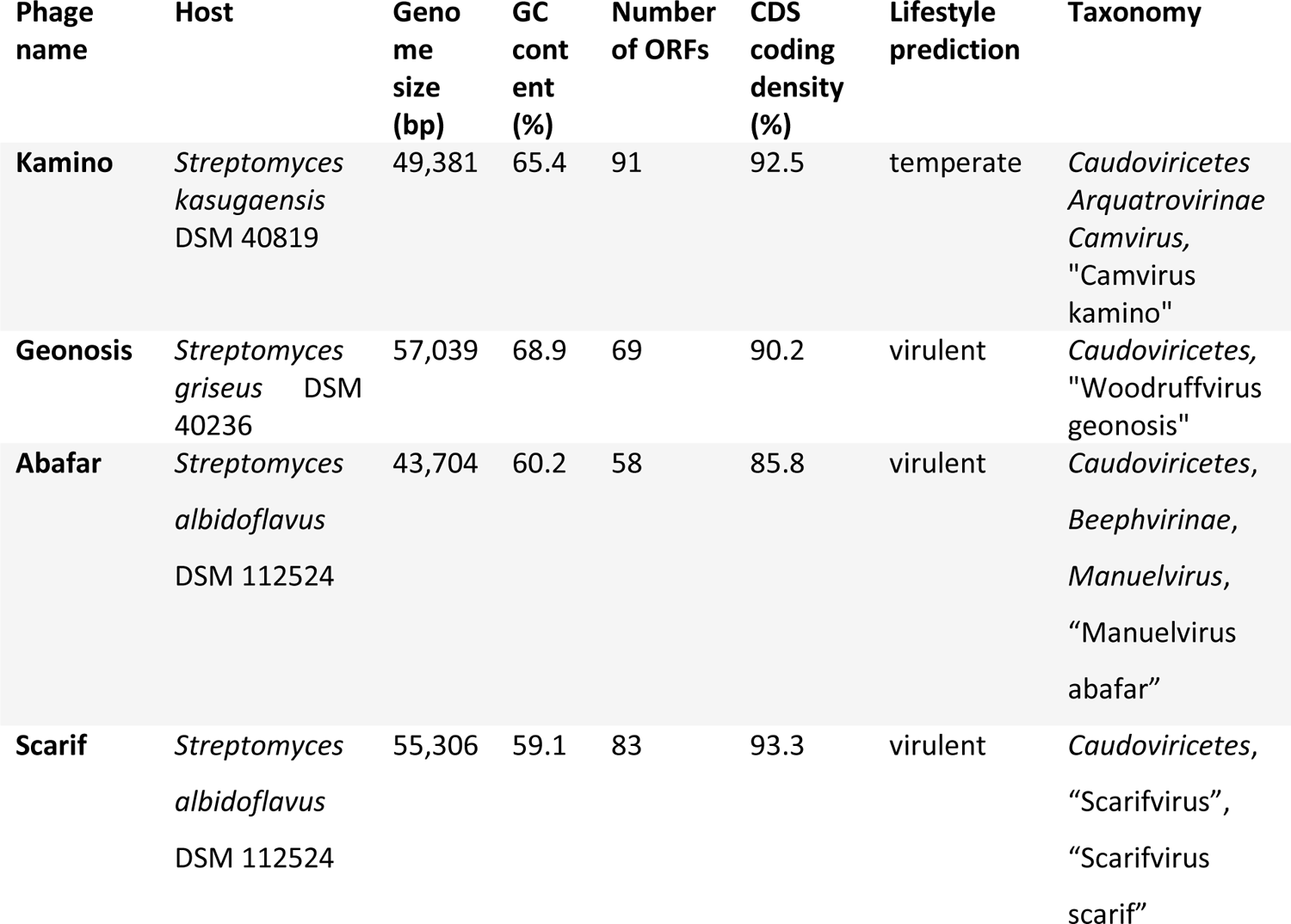
Summary of genomic features of the four *Streptomyces* phages.

The genome of phage Abafar consists of 43,704 bp (GC% 60.2) with 58 predicted open reading frames (ORF) and 15 genes for tRNAs (Figure 4). BLASTn analysis against NCBI database for viruses (taxid: 10239) identified four closely related phages, classified members of the genus *Manuelvirus*. All of them share a similar genome organization with functional gene clusters for packaging, and structural proteins with an embedded gene for a putative endolysin between the genes encoding the terminase large subunit and a portal protein (Figure S1). The cluster for replication contains conserved genes coding for a primase, a helicase and different nucleases. No genes related to lysogeny were identified, which is in line with the prediction of Abafar being virulent.

**Figure 4:**
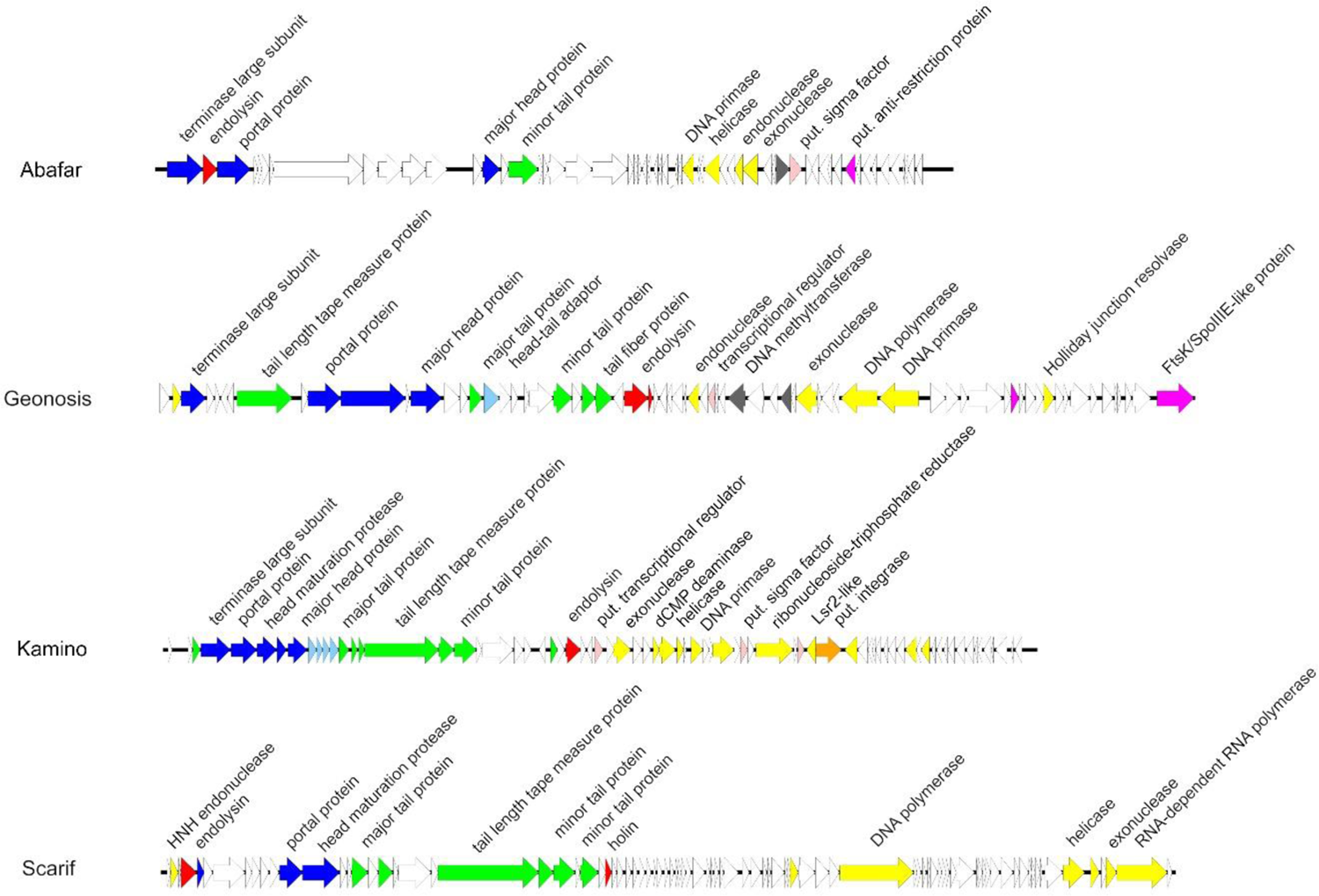
Annotated genomes of phages Abafar, Geonosis, Kamino and Scarif. Colouring is based on the PHROG colour code for functional clusters (Terzian et al., 2021) (orange: integration and excision; blue: head and packaging; purple: transcription; light blue: connector; green: tail; red: lysis; yellow: DNA, RNA and nucleotide metabolism; pink: moron, auxiliary metabolic gene and host takeover; dark grey: other).

Phage Geonosis has a genome size of 57,039 bp with an average GC content of 68.3 % comprising 69 ORFs but no tRNA genes (Figure 4). Comparison with other related Woodruffviruses like phage YDN12 ^32^ revealed a similar genome organization with gene clusters for packaging, structural head and tail proteins, lysis comprising endolysin and holin genes and replication containing characteristic genes for an endonuclease, a DNA methyltransferase, an exonuclease, a DNA polymerase and a DNA primase (Figure S2). Furthermore, phage Geonosis also harbours genes for a putative Holliday junction resolvase and an FtsK-like protein, respectively, that might also be used during replication.

Phage Kamino features a genome size of 49,381 bp with an average GC content of 65.4 % harbouring 91 ORFs, but no tRNA genes (Figure 4). BLASTn analysis identified related phages of the genus *Camvirus* like Endor1 ^18^, phiCAM ^33^, Verabelle or Vanseggelen ^34^. All of them share the same genome organization with characteristic gene clusters for structure, replication and lysis (Figure S3). In contrast to the other three isolated phages, phage Kamino harbors a gene for a serine integrase (Kamino_00049) and additional genes putatively involved in transcriptional regulation (Kamino_00029, Kamino_00043 and Kamino_00046) which is in line with the AI prediction of a temperate lifestyle.

Genomic analysis of phage Scarif revealed a genome size of 55,306 bp (GC%: 59.1) with 89 predicted ORFs organized in characteristic clusters for replication including genes for a DNA polymerase, a helicase, an exonuclease or an RNA-dependent RNA polymerase and for head and tail structure comprising genes for a portal protein, minor and major tail proteins and a tail length tape measure protein (Figure 4). Given that this protein determines the tail length of the phage and assuming a tail length of 1.5 Å per amino acid residue, the calculated length (∼274 nm) fits to the size measured via electron microscopy ^35^. Generally, phage Scarif shares this genomic organization with phages belonging to the genus *Rimavirus* (Figure S4).

VIRIDIC analysis (Suppl. Table,^36^) identified phages Abafar, Geonosis and Kamino as putative new species within the genera *Manuelvirus*, *Woodruffvirus* and *Camvirus*, respectively, according to the ICTV rules with 95 % and 70 % nucleotide sequence identity over the length of the genome as species and genus demarcation criteria, respectively ^37,38^. In contrast, based on this analysis, phage Scarif forms a new genus that we suggest to call “Scarifvirus”.

### Sensitivity towards antiphage compounds

Recent studies have shown that some molecules produced by *Streptomyces* have pronounced impact on phage infection ^22,24^. To gain first insights, we tested the sensitivity of the novel phages against a set of known antiphage compounds, including the aminoglycoside antibiotics apramycin and hygromycin, as well as the DNA-intercalating molecules daunorubicin and actinomycin D (Figure S5)^14,15^. Therefore, the three broad host-range phages Kamino, Abafar and Scarif were tested on *S. albidoflavus* M145 as host, carrying resistance plasmids for apramycin (pIJLK04) and hygromycin (pIJLK01). The DNA-intercalating molecules daunorubicin and actinomycin D were tested on wild type strains. Geonosis was tested on its isolation host *S. griseus* as wild type on DNA-intercalating molecules and on *S. griseus* pIJLK04 on apramycin, since its narrow host range restricts the testing of sensitivity on other host organisms (Figure 5). For all four phages, the strongest inhibitory effect was observed when challenged with the DNA-intercalating molecule daunorubicin. At a concentration of 6 µM, daunorubicin decreased the phage titre of Kamino 10^4^-fold and for Abafar and Scarif 10^5^ to 10^6^-fold. Since *S. griseus* did not form a bacterial lawn on concentrations higher than 1 µM, we only observed a small decrease in phage titre of ∼70 %. *S. albidoflavus* M145 also showed considerable growth defects on elevated daunorubicin concentrations. In contrast, the DNA intercalating peptide antibiotic actinomycin D showed only minor effects on the phage infections compared to daunorubicin. At the highest concentration of 6 µM all phages infecting *S. albidoflavus* M145 showed a decrease in titre of ∼50-70%, whereas the titre of Geonosis remained stable. The broad host-range phages Kamino, Abafar and Scarif showed a decrease in titre of 10^1^ to 10^3^ on 25 µg/ml apramycin, while Geonosis remained stable in titre at the highest concentration tested. The lowest impact on infection of the phages was exerted by hygromycin, which seemed to have no effect on plaque formation of Kamino and Scarif and only little effect on the plaque formation of Abafar, decreasing the titre by ∼75 % at the highest concentration of 50 µg/ml.

**Figure 5:**
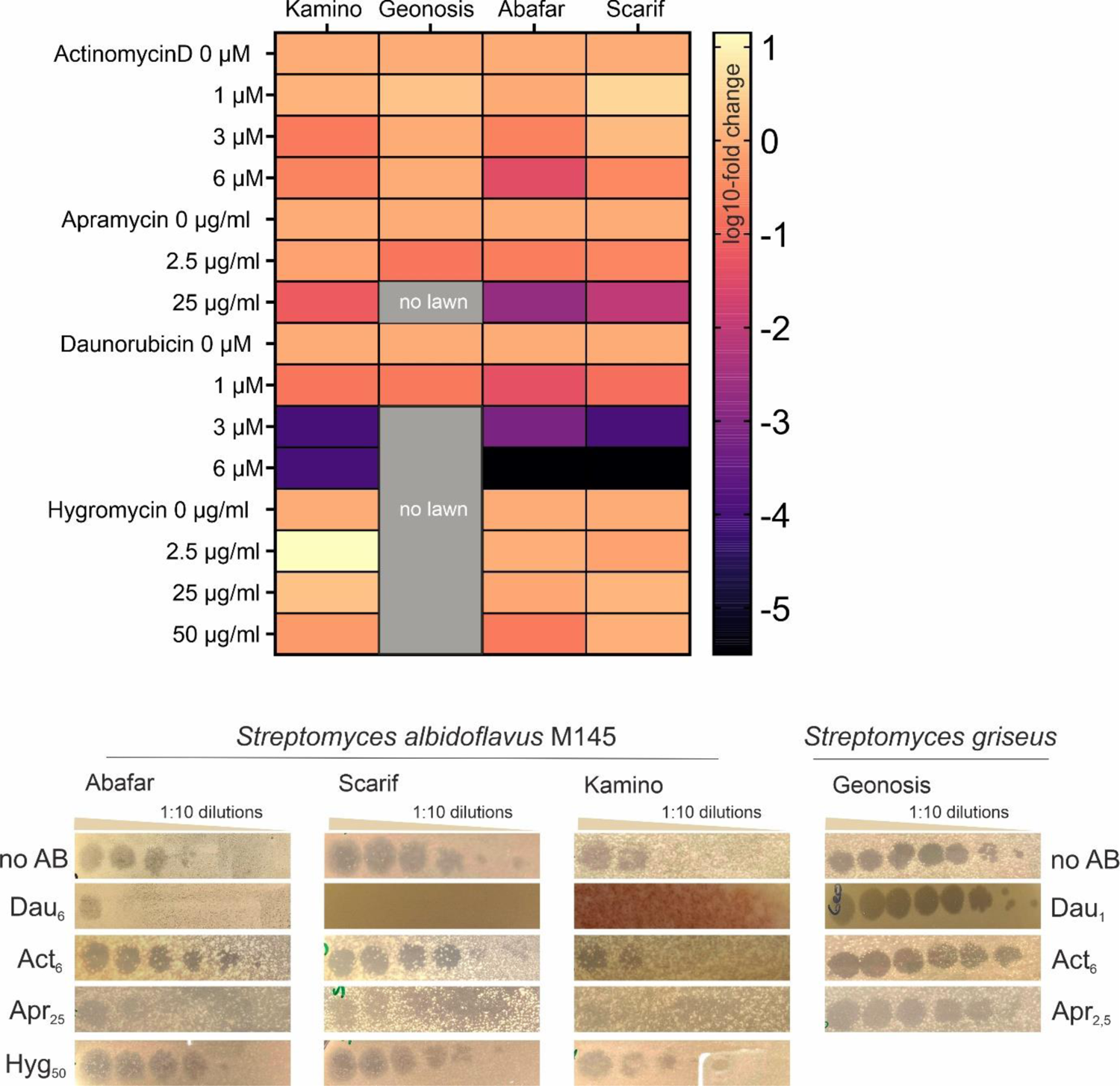
Screening of antiphage small molecules. On top is a heat map that represents the reaction of each phage (column) towards different concentrations of known anti-phage small molecule (row). The colour indicates the log10-fold change of phage titre. Below plaque images of the screening of each phage are shown exemplarily (n= 2 for each phage)

## Discussion

In this study, we present the isolation and characterization of four novel *Streptomyces* phages. Phage Kamino was isolated on *S. kasugaensis,* Geonosis was isolated on *S. griseus* and Abafar and Scarif were isolated on *S. albidoflavus.* Phage Kamino is predicted to be a temperate phage, carrying a serine integrase on its genome, whereas Geonosis, Abafar and Scarif are predicted to be strictly virulent. Three of the four phages described in this study have a broad host range with productive infections on 7 to 15 different *Streptomyces* species (out of 45 different strains tested).

Of the phages reported in this study, Kamino and Geonosis amplify well in liquid cultures (Figure 2). In particular, Kamino showed highly efficient infection of *S. kasugaensis* already at very low starting titres (10^2^ PFU/ml). In contrast to this, Abafar and Scarif show little amplification in liquid infections and no effect on the growth of their host organism *S. albidoflavus* M145. This is, in fact, not an unusual feature of phages infecting *Streptomyces* and has been reported previously ^34,39^.

When compared to the recently reported *Streptomyces* phages Vanseggelen and Verabelle, which also infect several different species ^34^ (Figure S4), two major differences were detected in the genome of phage Kamino ^40^. First, the gene for the integrase (Kamino_00049) shows no similarity at the nucleotide level to the integrase genes in Vanseggelen and Verabelle, secondly the gene annotated as a putative tail fibre in Verabelle and Vanseggelen, respectively, also reveals only weak similarities to its homolog in Kamino (Kamino_00020). However, the differences in their tail fibres compared to their otherwise conserved genome structure might explain their different host range behaviour as those structures are major players in virus-host interactions ^40^. Moreover, broad host range phages like Kamino, equipped with a serine integrase for a temperate lifestyle, can serve as powerful tools for genetic manipulation across various strains ^8,21^. In this context, novel integrative plasmids could be engineered utilizing the attP site of Kamino, expanding the repertoire of genetically modifiable *Streptomyces* strains.

Little is known so far about the characteristic phage traits of different morphotypes in phages infecting actinobacteria ^41^. In line with our findings, the majority of described *Streptomyces* phages belong to the morphotype of siphoviruses. Phage Abafar isolated in this study, however, might be a good example to study the differences in infection dynamics and adsorption of podoviruses. Genomic analysis of phage Abafar revealed that - in contrast to the three siphoviruses of this study - its genome contains 15 tRNA genes. Those phage-encoded tRNAs are considered to be used to evade host defense mechanisms that directly target tRNAs ^42,43^. Furthermore, we detected a gene (Abafar_00064) with an incomplete conserved domain that shows similarities to an ArdA-like anti-restriction protein. BLASTp analysis of the predicted amino acid sequence identified homologous proteins only in other podoviruses infecting *Streptomyces* species.

Recent years have seen an unprecedented expansion of our understanding of the prokaryotic immune system with more than 100 new systems been identified ^44^. It is a typical feature of immune systems that they confer protection against some, but certainly not against all types of viruses. This holds also true for antiphage molecules produced by *Streptomyces*, which were shown to affect phage infection to different extends ^14,15^. In this study, we observed the strongest antiphage effects for the DNA intercalating agent daunorubicin which inhibited all four tested phages. In contrast, the DNA-intercalating peptide antibiotic actinomycin D displayed only minor effects. These variations may arise from differences in their intercalation properties and/or from differences in molecule uptake. Significant differences between different phages and hosts were also observed for aminoglycoside antibiotics where so far only minor effects were observed for phages infecting *S. albidoflavus* (*S. coelicolor*) ^14^. For the phages described in this study, phage Abafar showed the highest sensitivity to all tested compounds, as a decrease in phage titre was observed for all tested molecules.

To further understand phage and host determinants conferring sensitivity to antiphage molecules the comparative analysis of a diverse set of phages is required. In fact, systematic phage collections proved highly valuable in assessing the efficacy of diverse defense systems against a wide spectrum of phages infecting a particular host or genus ^45^. Broad host range phages further provide the opportunity to determine context dependency by comparing the effect of a given defense system/antiphage molecule across different host species.

## Author Contributions

Conceptualization: B.R., C.R., J.W. J.F.; Data Curation, B.R., C.R.; Formal Analysis: B.R., C.R., I.M.; Funding acquisition: J.W., J.O., J.F.; Investigation: B.R., C.R., I.M.; Methodology: All; Project administration: J.W. and J.F.; Resources: J.W., J.O., J.F.; Supervision: J.W., J.O., J.F.; Validation: All; Visualization: B.R., C.R., J.W.; Writing—original draft: B.R., C.R., J.W., J.F.; Writing—review and editing: All. All authors have read and agreed to the published version of the manuscript.

## Funding

We thank the Deutsche Forschungsgemeinschaft (SPP 2330, project 464434020 and 465136285) for financial support.

## Acknowledgements

We would like to thank Vikas Sharma for his help with phage genome analysis and Stephanie Held for her help and support.

## Conflicts of interest

The authors declare no conflict of interest.

## Notes

### Competing Interest Statement

The authors have declared no competing interest.

