## Supplementary material for "Isolation and characterization of the new *Streptomyces* phages Kamino, Geonosis, Abafar and Scarif infecting a broad range of host species": TableS1 and S2; Figures S1-5

**Content:**

Table S1: Bacterial strains used for host range analysis and efficiency of plating

Table S2: Bacterial strains used for host range analysis.

Supplementary Figure 1

Supplementary Figure 2

Supplementary Figure 3

Supplementary Figure 4

Supplementary Figure 5

Table S3: VIRDIC analysis of phages (extra)

### Supplementary Information

Table S1: Bacterial strains used for host range analysis and efficiency of plating.

Bacterial strains in this table were used to perform detailed experiments in the sections describing phage isolation and propagation, infection dynamics, plaque development and phage host range assay with EOP.

| **Strains** | **Genotype** | **Reference** |
|---|---|---|
| *Streptomyces afghaniensis* | WT | DSM 40228 |
| *Streptomyces albaduncus* | WT | DSM 40478 |
| *Streptomyces albidoflavus* (host Abafar & Scarif*)* | WT | DSM 112524 |
| *Streptomyces albidoflavus* M145 | *S. coelicolor* A3(2) lacking plasmids SCP1 and SCP2 | (Bentley *et al.*, 2002) |
| *Streptomyces albidoflavus* M145 pIJLK01 | Carrying the integrative plasmid pIJLK01, $Hyg^R$ | (Kever et al., 2022) |
| *Streptomyces albidoflavus* M145 pIJLK04 | Carrying the integrative plasmid pIJLK04, $Apr^R$ | (Kever et al., 2022) |
| *Streptomyces albulus* | WT | DSM 40492 |
| *Streptomyces antibioticus* | WT | DSM 40234 |
| *Streptomyces avermitilis* | WT | DSM 46492 |
| *Streptomyces azureus* | WT | DSM 40106 |
| *Streptomyces celluloflavus* | WT | DSM 40839 |
| *Streptomyces chartreusis* | WT | DSM 40085 |
| *Streptomyces clavuligerus* | WT | DSM 738 |
| *Streptomyces fimbriatus* | WT | DSM 40942 |
| *Streptomyces fradiae* | WT | DSM 40063 |
| *Streptomyces galbus* | WT | DSM 40089 |
| *Streptomyces griseofuscus* | WT | DSM 40191 |
| *Streptomyces griseorubens* | WT | DSM 40160 |
| *Streptomyces griseus* (host Geonosis) | WT | DSM 40236 |
| *Streptomyces giseus* pIJLK04 | Carrying the integrative plasmid pIJLK04, $Apr^R$ | This study |
| *Streptomyces humidus* | WT | DSM 40263 |
| *Streptomyces hygroscopicus* | WT | DSM 40578 |

| | | |
|---|---|---|
| *Streptomyces inusitatus* | WT | DSM 41441 |
| *Streptomyces kanamyceticus* | WT | DSM 40500 |
| *Streptomyces kasugaensis* (host Kamino) | WT | DSM 40819 |
| *Streptomyces lavendulae* | WT | DSM 40069 |
| *Streptomyces longispororuber* | WT | DSM 40599 |
| *Streptomyces luridus* | WT | DSM 40081 |
| *Streptomyces niveus* | WT | DSM 40088 |
| *Streptomyces nodosus* | WT | DSM 40109 |
| *Streptomyces olivaceus* | WT | DSM 41536 |
| *Streptomyces purpurascens* | WT | DSM 40310 |
| *Streptomyces rimosus* | WT | DSM 40260 |
| *Streptomyces sulfonofaciens* | WT | DSM 41679 |
| *Streptomyces thioluteus* | WT | DSM 40027 |
| *Streptomyces venezuelae* NRRL B-65442 | WT | DSM 112328 |
| *Streptomyces violaceus* | WT | DSM 40082 |
| Streptomyces *viridosporus* | WT | DSM 40243 |

Table S2: Bacterial strains used for host range analysis.

Table S2 includes all *Streptomyces* strains which were used for the host range assay. Strains which showed productive infection were further tested and are mentioned in Table S1 in detail

| ***Streptomyces* strains used for the host range** |
|---|
| *Streptomyces afghaniensis* |
| *Streptomyces albaduncus* |
| *Streptomyces albidoflavus* |
| *Streptomyces alboniger* |
| *Streptomyces albulus* |
| *Streptomyces anandii* |
| *Streptomyces antibioticus* |
| *Streptomyces avermitilis* |
| *Streptomyces avidinii* |
| *Streptomyces azureus* |
| *Streptomyces bluensis* |

| |
|---|
| *Streptomyces celluloflavus* |
| *Streptomyces chartreusis* |
| *Streptomyces chrestomyceticus* |
| *Streptomyces clavuligerus* |
| *Streptomyces echinatus* |
| *Streptomyces eurocidicus* |
| *Streptomyces fradiae* |
| *Streptomyces fimbriatus* |
| *Streptomyces galbus* |
| *Streptomyces griseus* |
| *Streptomyces griseofuscus* |
| *Streptomyces griseorubens* |
| *Streptomyces humidus* |
| *Streptomyces inusitatus* |
| *Streptomyces kanamyceticus* |
| *Streptomyces kasugaensis* |
| *Streptomyces lavendulae* |
| *Streptomyces litmocidini* |
| *Streptomyces longispororuber* |
| *Streptomyces luridus* |
| *Streptomyces mutomycini* |
| *Streptomyces niveus* |
| *Streptomyces nodosus* |
| *Streptomyces olivaceus* |
| *Streptomyces purpurascens* |
| *Streptomyces rapamycinicus* |
| *Streptomyces rimosus* |
| *Streptomyces scabiei* |
| *Streptomyces sulfonofaciens* |
| *Streptomyces thioluteus* |
| *Streptomyces venezuelae* |
| *Streptomyces violaceus* |
| *Streptomyces viridosporus* |

*Streptomyces wellingtoniae*

### 39 Supplementary Figure 1

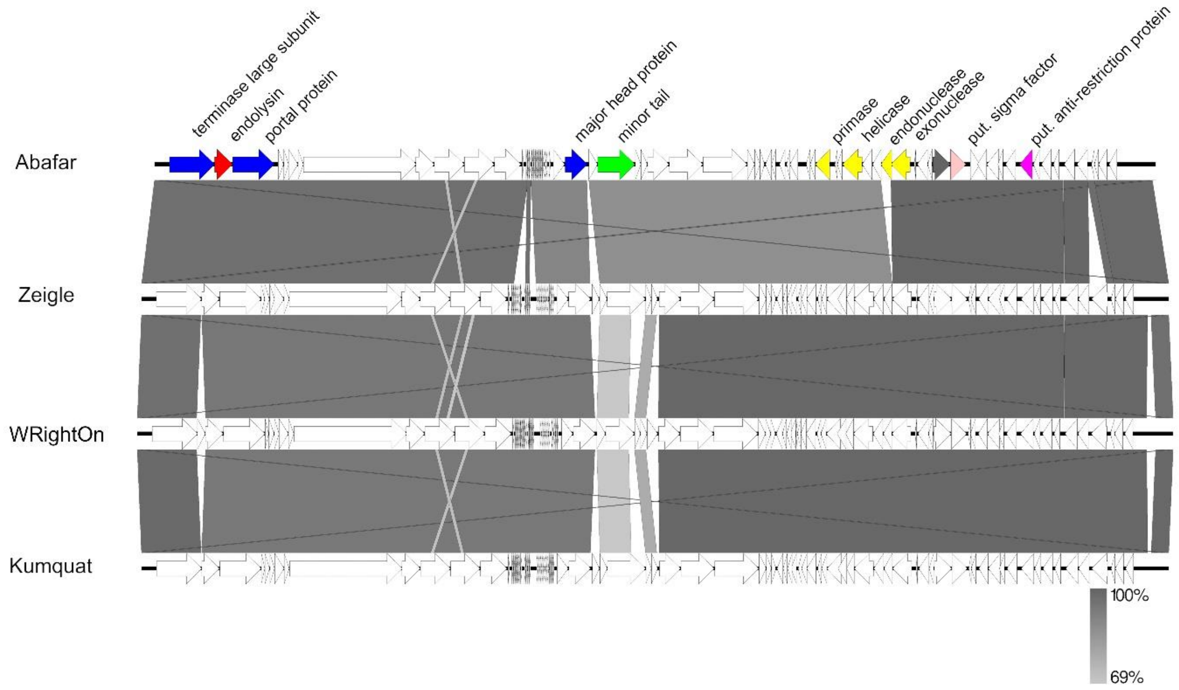


Figure S1: Synteny plot of phage Abafar compared to other manuelviruses at the nucleotide level. The figure was generated with EasyFig (Sullivan et al., 2011). Colouring is based on the PHROG colour code for functional clusters (Terzian et al., 2021) (orange: integration and excision; blue: head and packaging; purple: transcription; light blue: connector; green: tail; red: lysis; yellow: DNA, RNA and nucleotide metabolism; pink: moron, auxiliary metabolic gene and host takeover; dark grey: other).

### 40 Supplementary Figure 2

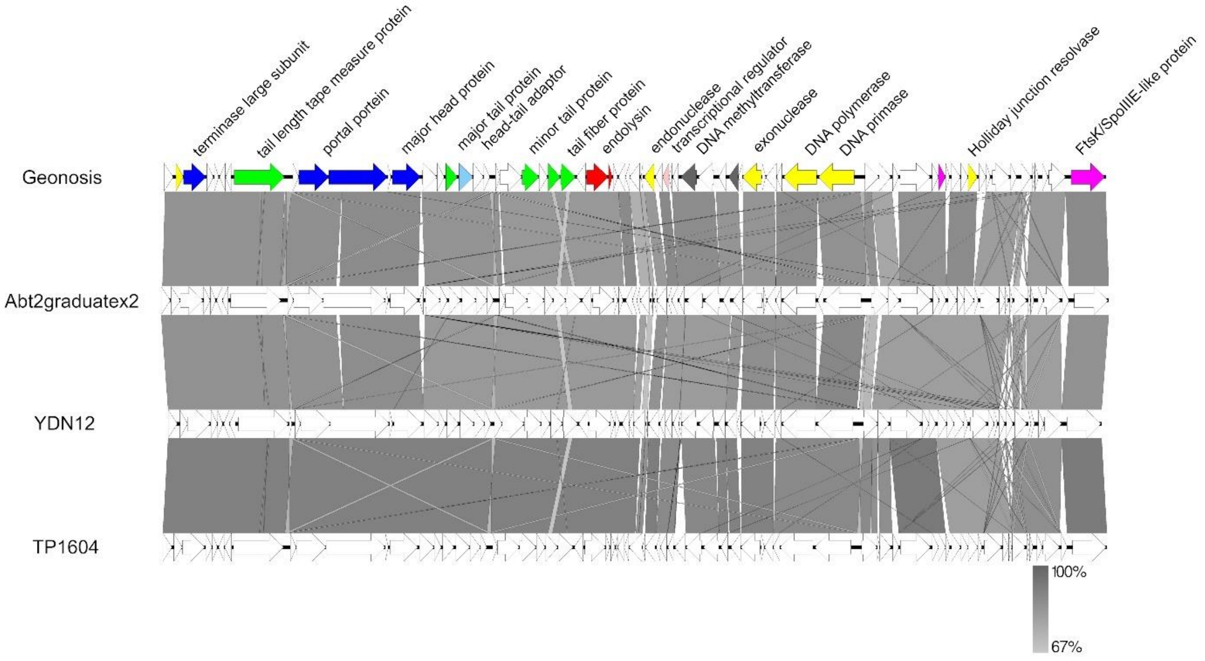


Figure S2: Synteny plot of phage Geonosis compared to other woodruffviruses at the nucleotide level. The figure was generated with EasyFig (Sullivan et al., 2011). Colouring is based on the PHROG colour code for functional clusters (Terzian et al., 2021) (orange: integration and excision; blue: head and packaging; purple: transcription; light blue: connector; green: tail; red: lysis; yellow: DNA, RNA and nucleotide metabolism; pink: moron, auxiliary metabolic gene and host takeover; dark grey: other).

Supplementary Figure 3

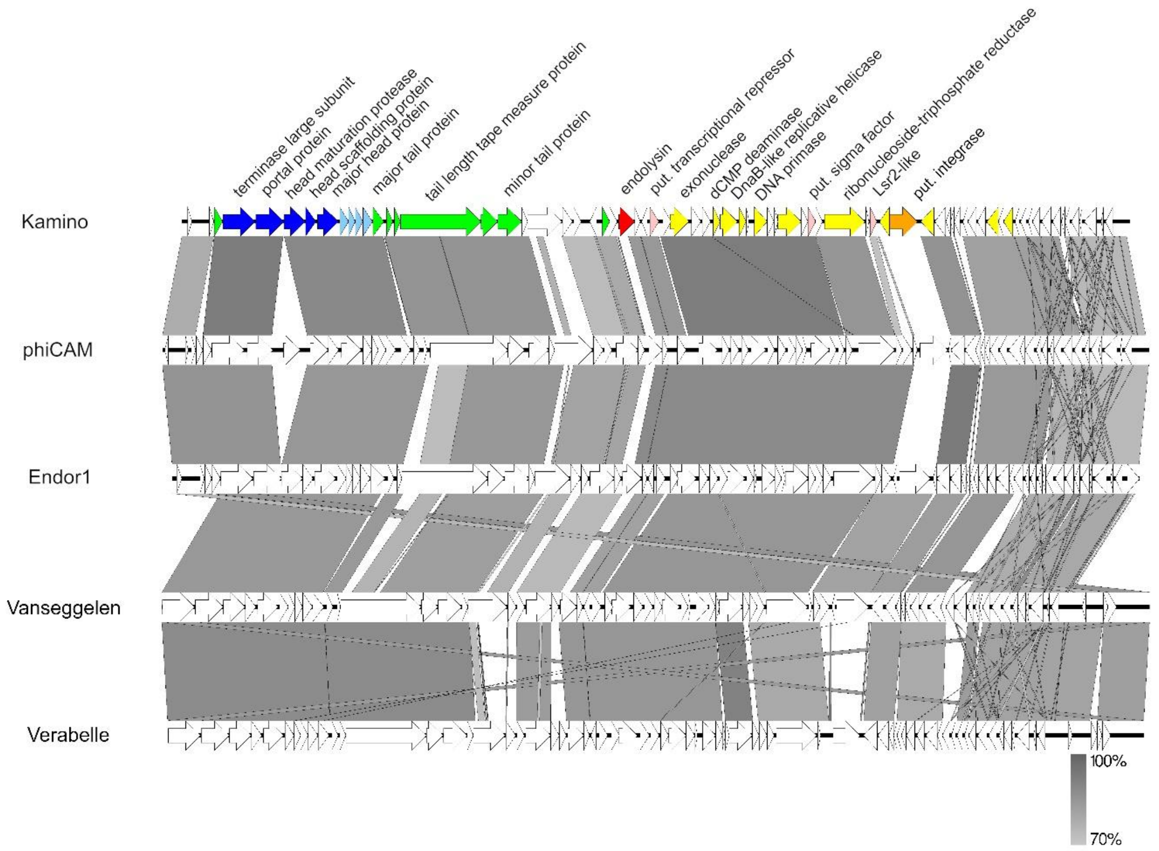


Figure S3: Synteny plot of phage Kamino compared to other camviruses at the nucleotide level. The figure was generated with EasyFig (Sullivan et al., 2011). Colouring is based on the PHROG colour code for functional clusters (Terzian et al., 2021) (orange: integration and excision; blue: head and packaging; purple: transcription; light blue: connector; green: tail; red: lysis; yellow: DNA, RNA and nucleotide metabolism; pink: moron, auxiliary metabolic gene and host takeover; dark grey: other).

44 Supplementary Figure 4

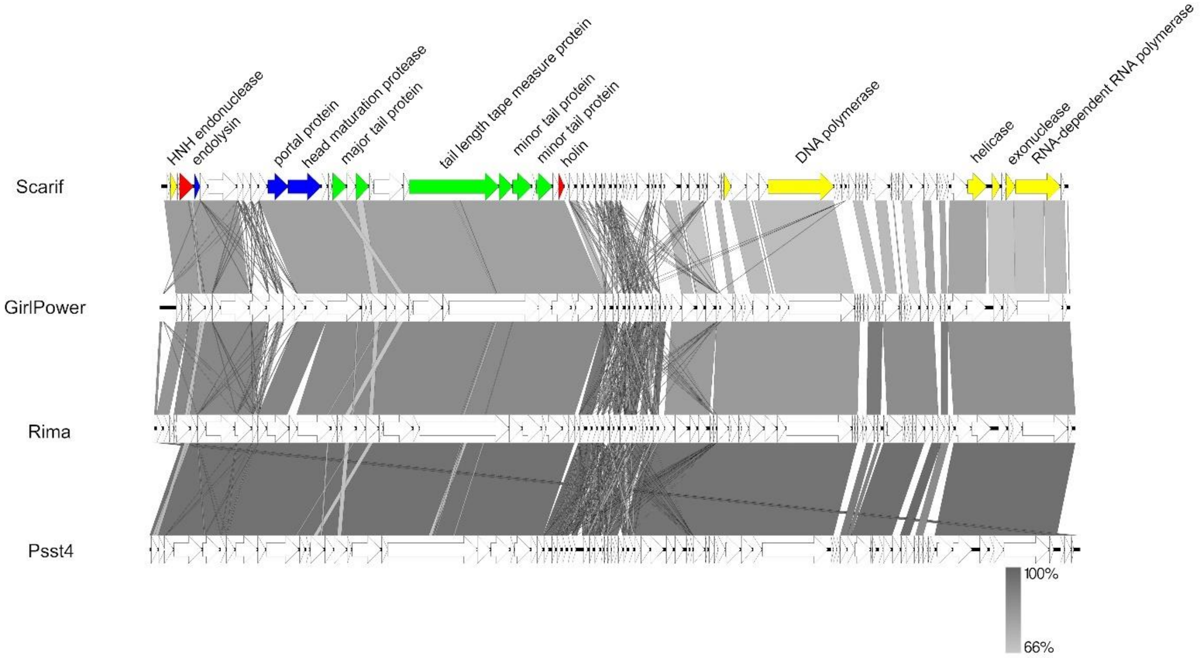


Figure S4: Synteny plot of phage Scarif compared to closely related members of Rimavirus at the nucleotide level. The figure was generated with EasyFig (Sullivan et al., 2011). Colouring is based on the PHROG colour code for functional clusters (Terzian et al., 2021) (orange: integration and excision; blue: head and packaging; purple: transcription; light blue: connector; green: tail; red: lysis; yellow: DNA, RNA and nucleotide metabolism; pink: moron, auxiliary metabolic gene and host takeover; dark grey: other).

45

Supplementary Figure 5:

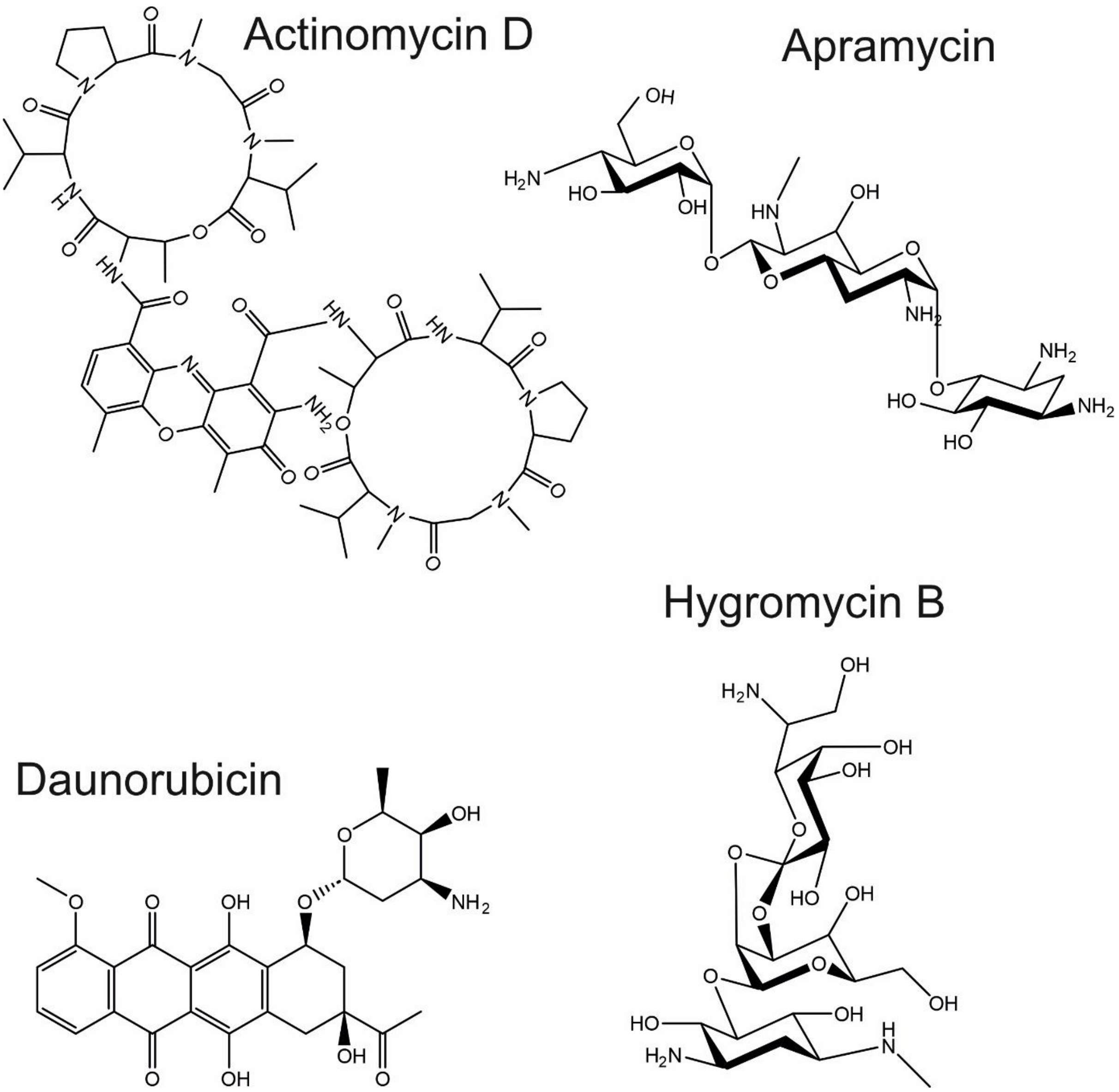


Figure S5: Molecular structure of antiphage compounds used in this study. Displayed are the aminoglycoside antibiotics apramycin and hygromycin as well as the DNA-intercalating anthracyclines daunorubicin and the peptide antibiotic actinomycin D.
